# Finding Haplotypic Signatures in Proteins

**DOI:** 10.1101/2022.11.21.517096

**Authors:** Jakub Vašíček, Dafni Skiadopoulou, Ksenia G. Kuznetsova, Bo Wen, Stefan Johansson, Pål R. Njølstad, Stefan Bruckner, Lukas Käll, Marc Vaudel

## Abstract

The non-random distribution of alleles of common genomic variants produces haplotypes, which are fundamental in medical and population genetic studies. Consequently, protein-coding genes with different co-occurring sets of alleles can encode different amino acid sequences: protein haplotypes. These protein haplotypes are present in biological samples, and detectable by mass spectrometry, but are not accounted for in proteomic searches. Consequently, the impact of haplotypic variation on the results of proteomic searches, and the discoverability of peptides specific to haplotypes remain unknown. Here, we study how common genetic haplotypes influence the proteomic search space and investigate the possibility to match peptides containing multiple amino acid substitutions to a publicly available data set of mass spectra. We found that for 9.96 % of the discoverable amino acid substitutions encoded by common haplotypes, two or more substitutions may co-occur in the same peptide after tryptic digestion of the protein haplotypes. We identified 342 spectra that matched to such multi-variant peptides, and out of the 4,251 amino acid substitutions identified, 6.63 % were covered by multi-variant peptides. However, the evaluation of the reliability of these matches remains challenging, suggesting that refined error rate estimation procedures are needed for such complex proteomic searches. As these become available and the ability to analyze protein haplotypes increases, we anticipate that proteomics will provide new information on the consequences of common variation, across tissues and time.

## Introduction

Linkage disequilibrium (LD) describes the non-random correlation between alleles at different positions in the genome in a population. LD arises when alleles at nearby sites co-occur on the same haplotype more often than expected by chance. When haplotypes are located in protein-coding portions of the genome, they can alter protein sequences, forming so-called *protein haplotypes*, as defined by Spooner *et al*. (1). Based on the co-occurence of alleles in the 1000 Genomes Project (2) and their *in silico* translation, Spooner et al. (1) created a list of possible protein haplotype sequences. Notably, they stress that for one in seven genes, the most frequent protein haplotype differs from the reference sequence in Ensembl (3). In precision medicine, probing the proteotype – the actual state of the proteome – adds valuable information concerning the relationship between the genotype and the phenotype (4). Therefore, it is important that genetic information including LD is taken into account in proteomics searches.

Proteins in biological samples can be identified by liquid chromatography coupled to mass spectrometry (LC-MS), usually after digestion into peptides (5). Then, the measured spectra are matched to a database of expected protein sequences using a search engine (6). The identified peptides are used to infer the presence of proteins (7) along with potential post-translational modifications (PTMs) (8). When the peptides cover the relevant parts of the protein sequences, it is also possible to discover the product of alternative splicing or genetic variation (9). In precision medicine, proteomic searches need to be adapted to individual patient profiles by extending the search space to include non-canonical sequences (10).

This challenge is addressed by proteogenomics – the scientific field integrating genomics and proteomics into a joint approach (9,11). Recent work, mainly in the domain of cancer research, has shown that accounting for genetic variation in proteomic analyses provides the means to discover non-canonical proteins. Umer *et al*. (12) have developed a tool to generate databases of variant proteins derived from single nucleotide polymorphisms (SNPs), insertions and deletions, and the three-frame translation of pseudogenes and non-canonical transcripts, appended with a database of canonical proteins (12). Levitsky *et al*. (13) use measures of proteome coverage including variant peptides to verify the presence of single amino acid variants. Choong *et al*. (14) proposed an algorithm to generate the optimal number of protein sequences containing combinations of amino acid substitutions possibly occurring in the same tryptic peptide. In their approach, the database includes not only the combinations of alleles encoded by haplotypes, but all combinations possible per peptide. Lobas *et al*. (15,16) showed that peptides containing variation were 2.5 to 3 times less likely to be identified than canonical peptides. Wang *et al*. (17) have analyzed data for 29 paired healthy human tissues from the Human Proteome Atlas project to detect amino acid variants at the protein level. However, the majority of amino acid variants predicted from exome sequencing could not be detected (17), suggesting that proteogenomics remains highly challenging and methods for discovering non-canonical proteins need further development.

Here, we used the protein haplotypes generated by Spooner et al. (1) to evaluate the ability of mass spectrometry-based proteomics to identify peptides encoded by combinations of variants in LD. We show that in some protein haplotypes, multiple amino acid substitutions affect the same peptide after digestion. Those protein haplotypes can only be identified if the combinations of amino acid variants are included in the search space, and several of these protein haplotypes are predicted to be more common than the reference sequence. Then, we mined the publicly available data from Wang et al. (17) for peptides including a combination of amino acid variants, demonstrating how such peptides can be identified according to the standards of the field, but also how the quality control of the results remains challenging.

## Results

### The consequence of haplotypes on the proteomics search space

We digested *in silico* the protein sequences translated from haplotypes obtained from Spooner *et al*. (1) using the canonical cleavage pattern of trypsin, allowing for up to two missed cleavages. Note that indels were not considered, and we focused only on common variants with a minor allele frequency > 1 % in any population of the 1000 Genomes Project (2), see methods for details. After excluding contaminants, this yielded 2,470,700 unique tryptic peptide sequences of length between 8 and 40 amino acids (Figure 1A). The coverage of protein sequences from Ensembl can be partitioned as follows: 80.04 % can only be covered by canonical peptides, 7.59 % map to peptides that may contain one or multiple amino acid substitutions, and the remaining yields sequences that are either too short or too long to be identified. The vast majority of peptides discoverable in proteomic studies therefore maps to canonical sequences, making it challenging for non-targeted approaches to assess the allelic status of a common genetic variant using proteomics, in agreement with (15,16).

**Figure 1:**
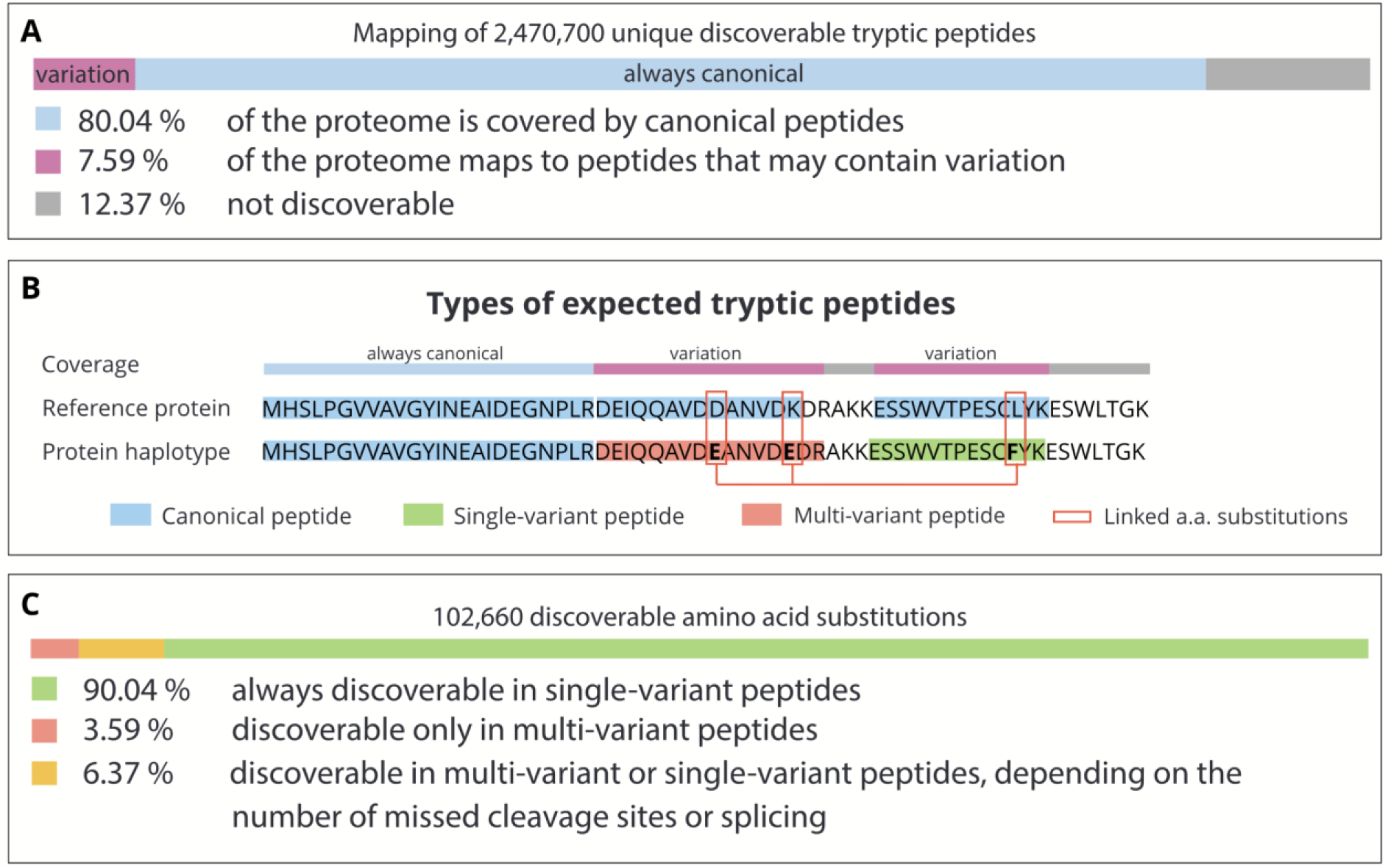
A: Proteome coverage expressed in terms of the percentage of amino acids, i.e., if 7 out of 100 residues belong to at least one discoverable peptide containing the product of a substitution, we say that 7 % of the proteome maps to peptides containing variation. See main text for details and methods for the handling of shared peptides. B: Example of a reference sequence aligned to another haplotype. The classes of peptides following the cleavage pattern of trypsin are highlighted by a colored background. Three amino acid substitutions encoded by this haplotype are marked by red rectangles. The “coverage” layer indicates the alignment applied to obtain numbers shown in section A. C: Distribution of variation in discoverable peptides. Amino acid variants are stratified based on the category of peptide in which the substitution caused by the respective variant can be identified.

We classify the obtained peptide sequences in three types (Figure 1B): (i) *canonical*, no haplotype is known to yield an amino acid substitution in the sequence of this peptide; (ii) *single-variant*, a haplotype encodes an amino acid substitution in the sequence of this peptide; and (iii) *multi-variant*, a haplotype encodes a set of two or more amino acid substitutions in the sequence of this peptide. In total, common haplotypes encode 102,660 amino acid substitutions, 90.04 % of them are found only in single-variant peptides, 3.59 % in multi-variant peptides, and 6.37 % in either single- or multi-variant peptides depending on the number of missed cleavages (Figure 1C). Note that substitutions in different isoforms of the same protein are reported separately by Spooner et al. (1), creating multiple consequences for the same genetic variant. The total number of amino acid substitutions is consequently higher than the number of genetic variants. Interestingly, based on the frequencies among all participants in the 1000 Genomes project, 21 % and 24.3 % of the amino acid substitutions discoverable in single-variant and multi-variant peptides, respectively, occur in protein haplotypes that are predicted to be more frequent than the Ensembl reference sequence. If these alleles are not accounted for, proteomics analyses will therefore not be able to identify these parts of the genome for the majority of individuals.

Peptides can be classified based on their ability to distinguish between protein sequences. We propose the following categories: (i) *non-specific* peptides map to the products of different genes; (ii) *protein-specific* peptides map to multiple sequences, which are all products of the same gene; and (iii) *proteoform-specific* peptides map uniquely to a single form of a protein (i.e., single splice variant and haplotype), referred to as proteoform (18). In this classification, based on the identification of a proteoform-specific peptide, one can uniquely identify products of a given gene. A protein-specific peptide allows for discriminating certain groups of proteoforms, but does not yield a single candidate sequence (e.g., it determines which amino acid substitution is present, but maps to multiple splicing variants). Non-specific peptide sequences map to multiple genes, where the sequence of one gene matches the sequence of another, making it challenging to infer which protein is covered. We found 182,821 distinct non-specific peptide sequences, covering up to 14.58 % of the proteome. The prevalence of canonical, single-variant, and multi-variant peptides among the above introduced types is displayed in Figure 2. As expected intuitively, peptides containing the product of one or multiple variants present a higher ability to distinguish between protein products of different genes, and between proteoforms of the same gene.

**Figure 2:**
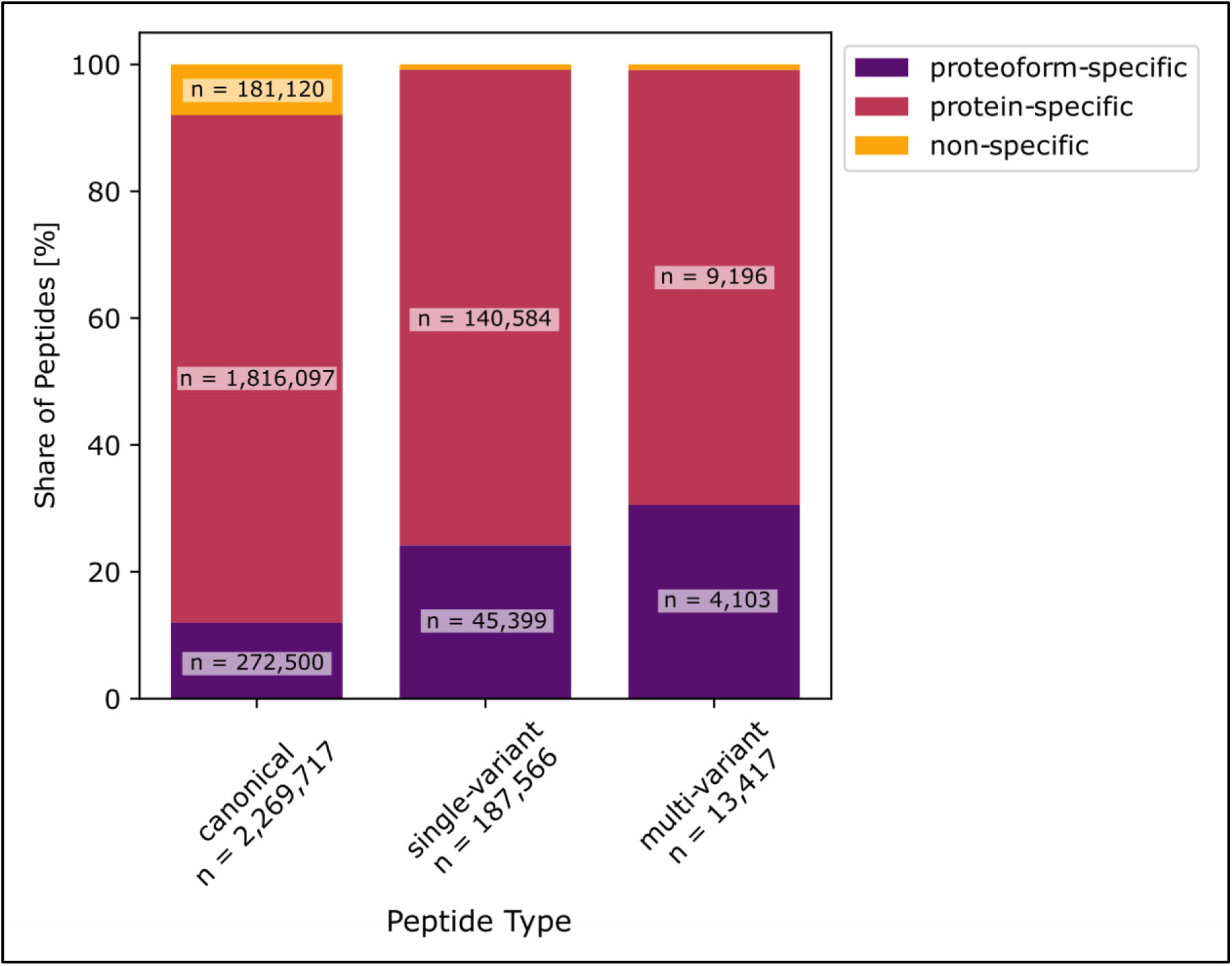
Classification of peptides based on their ability to distinguish between protein sequences (bar color) and to identify amino acid substitutions (position on X axis). The height of the bars represent the distribution of categories (non-specific, protein-specific, proteoform-specific) among the peptide types (canonical, single-variant, multi-variant).

### Matching multi-variant peptides to mass spectra

To investigate the prevalence of spectra matching multi-variant peptides encoded by common haplotypes, and the quality of the obtained matches, we searched the deep proteomics data of healthy tonsil tissue made available by Wang *et al*. (17) against the sequences of common protein haplotypes using X!Tandem (19) as search engine without refinement procedure, and Percolator (20) with standard features for the evaluation of the confidence in all peptide to spectrum matches (PSMs). The resulting PSMs were thresholded at 1 % PSM-level false discovery rate (FDR). Note that our study focuses on evaluating the quality of the spectrum matches, a PSM-level FDR was therefore preferred to peptide-level statistics. After thresholding, 1,318,150 target PSMs remained (13,466 decoy PSMs would have passed the threshold), representing 176,193 unique peptide sequences (8,047 decoy peptide sequences would have passed the threshold), covering the alternative amino acid of 4,251 substitutions. The distribution of alternative alleles among single and multi-variant peptides (Figure 3A) mirrored the values obtained from the *in silico* digestion of protein haplotypes (Figure 1C). On average, the product of 2,095 substitutions were found per sample (2,194, 2,087, and 2,005 in sample 1, 2, and 3, respectively). The matched peptide sequences cover 21.75 % of the proteome predicted to map exclusively to canonical peptides, and 16.99 % of the proteome possibly mapping to peptides with substitutions (Figure 3B). Note however that 19,678 peptide sequences (identified in 231,181 PSMs) map to the products of multiple genes that cannot be distinguished, hence affecting coverage estimates.

**Figure 3:**
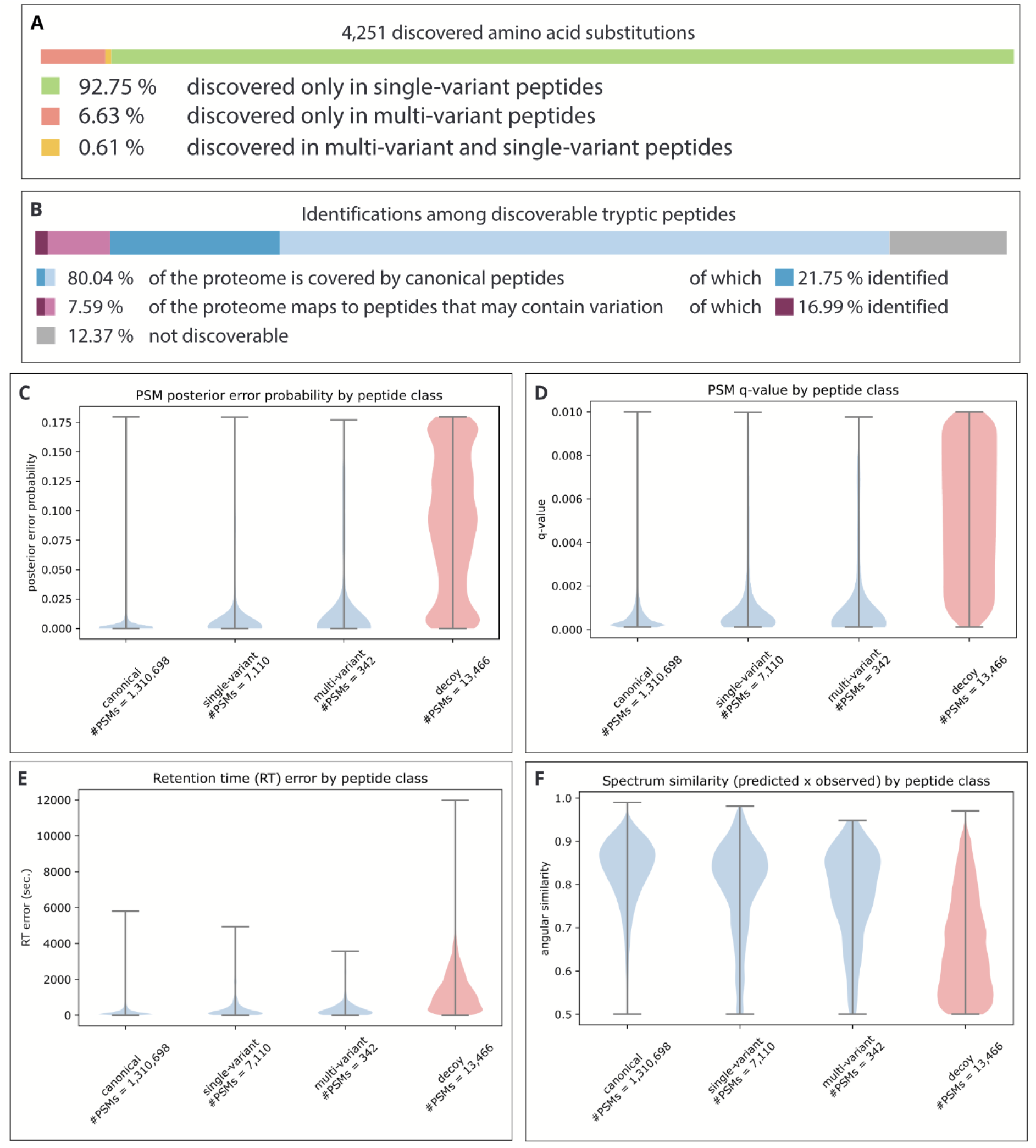
A: Coverage of the proteome by identified peptides, stratified by the possibility to contain variation. Lighter shades indicate the coverage by predicted peptides, darker shades represent the actual coverage by identified peptides. B: Distribution of variation in identified peptides. Amino acid variants are stratified based on the category of peptide in which the substitution caused by the respective variant can be identified. C-F: Distribution of four confidence measures among PSMs for peptide categories: posterior error probability (PEP), q-value, difference between observed and predicted retention time, and angular similarity between the observed and predicted spectrum. Decoy PSMs for this comparison were thresholded to 1% PSM-level FDR.

Out of the 1,318,150 spectra matched to peptides, 0.54 % were matched to single-variant peptides and 0.03 % were matched to multi-variant peptides. The share of spectra matched to variant peptides is thus lower than the expected error rate, and currently no method allows the evaluation of error rates in these subgroups of matches specifically. We thus investigated whether these classes of peptides showed signs of an overrepresentation of false positive matches. No substantial difference was noticeable in the density of the posterior error probabilities (PEPs) and q-values for all three classes of PSMs (Figure 3C and 3D), indicating that a more stringent FDR threshold would not alter the prevalence of variant peptides. We also compared the observed peptide retention time and fragmentation with predictions from DeepLC (21) and MS2PIP (22), respectively. Overall, the density of the distance to prediction in both retention time and fragmentation was very similar for all three classes of peptides (Figure 3E and 3F), displaying no obvious shift in the distribution which would have been indicative of a strong overrepresentation of false positives. Yet the distributions of variant and multi-variant peptides showed stronger tails towards high distance to prediction compared to non-variant peptides, indicative of the presence of false positive matches. In comparison, the distance to prediction for decoys showed high retention time difference and low spectrum similarity.

Quality metrics on all matches are available as supplementary material. Three examples, sampled from the multi-variant matches passing the FDR threshold at low, medium, and high PEP, representing high, medium, and low confidence, respectively, are displayed in Figure 4 along with the predicted spectra. As expected, the share of peaks matching predicted fragment ions decreases as the PEP increases: (A) the highly confident match presents an excellent coverage of the spectrum with fragment ion masses, with an extensive mapping of the peptide y-ion series, the retention time distance to prediction of 320.8 s represents only a fraction of the gradient (approx. 2.5 h), and the spectrum similarity to prediction, 0.79, shows good but not perfect agreement, which is in the lower range of the distribution of similarity scores for the canonical matches; (B) the medium confidence match presents a good coverage of the spectrum lacking prediction for many peaks, the agreement scores with retention time and fragmentation predictions are excellent; (C) the low confidence match presents a poor coverage of the spectrum with poor agreement with retention time and fragmentation predictions. In addition to passing commonly accepted statistical thresholds, the matches in Figure 4A and 4B would pass expert quality control. On the other hand, the match in 4C is most probably a false positive. Together, while these three sampled PSMs represent only a limited set of examples, they are very representative of the difficulty to confidently assess the presence of individual peptides from large proteomic experiments. This task is however important given that chimeric spectrum matches (23–26) and partial matches are known to be difficult to account for in error rate estimation (27,28).

**Figure 4:**
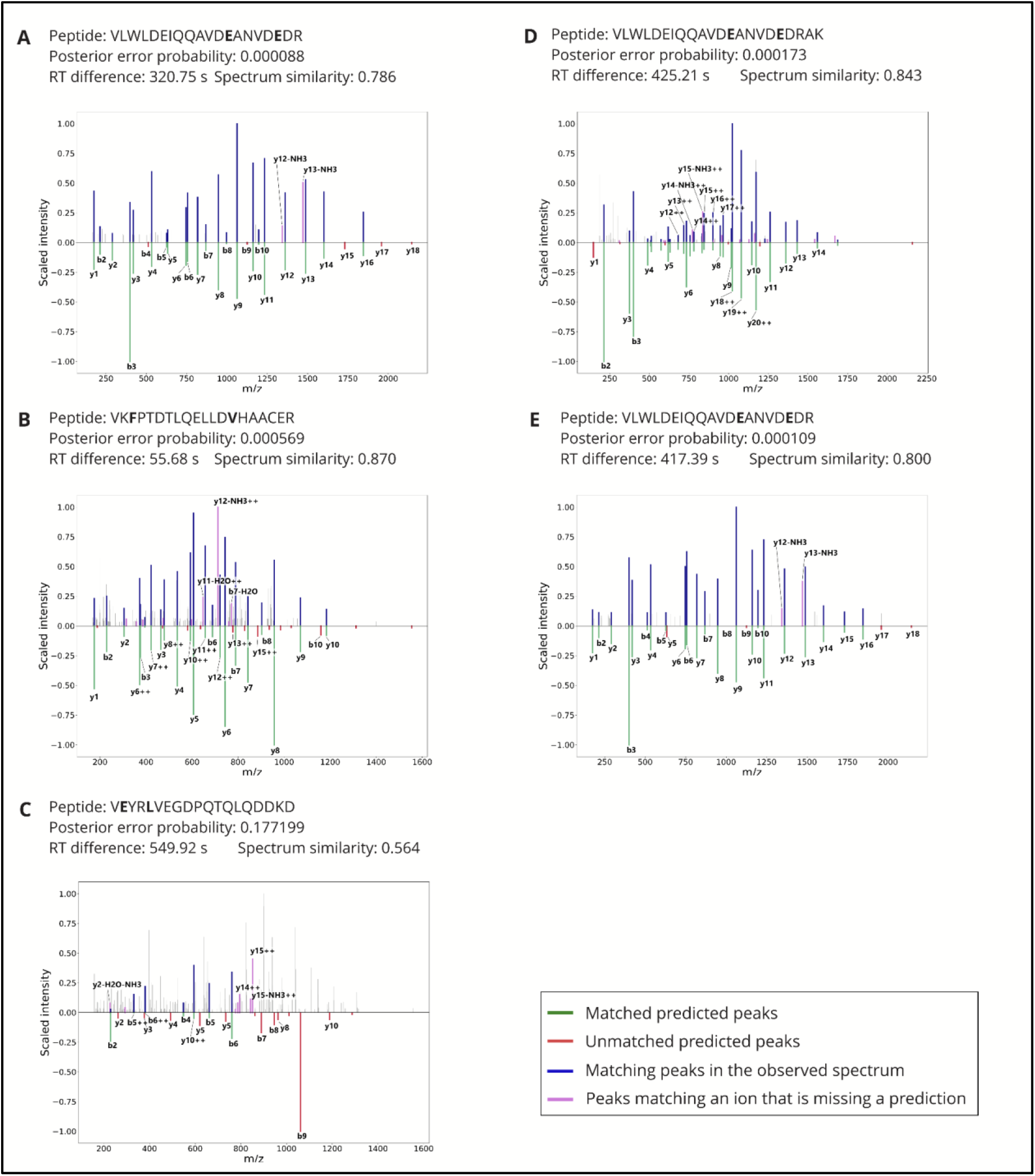
Quality control metrics and spectra of five multi-variant (PSMs). Amino acid substitutions are marked in bold. PSM A is among the 10 % top-scoring matches to multi-variant peptides by posterior error probability, B scores as the median value, and C is the lowest-scoring match to a multi-variant peptide. PSMs D and E are within the 5 top-scoring matches for the most common haplotype of IQGAP2. The posterior error probability as obtained from Percolator is listed along with retention time difference to prediction as obtained from DeepLC, and spectrum similarity with prediction as obtained from MS2PIP. The intensity of the measured spectrum is plotted (top; blue, pink, and gray) with the scaled predicted intensity mirrored (bottom; green and red). Peaks in the measured spectrum matching predictions are highlighted in blue, measured peaks matching an ion with a missing intensity prediction are highlighted in pink, while other measured peaks are plotted in gray. Note that in this representation, peaks matching a fragment ion with a predicted intensity of zero will not be annotated.

As highlighted by Spooner et al. (1), depending on the population studied, specific haplotypes often have higher frequencies than the canonical haplotype by Ensembl. For example, there are five haplotypes of the *IQ motif containing GTPase activating protein 2* (IQGAP2, ENSP00000274364) gene that have higher predicted frequencies than the canonical haplotype in the European population (with combined frequency of 84.9 % according to Spooner et al. (1)). These haplotypes encode a tryptic peptide containing two amino acid substitutions when compared to the canonical sequence in Ensembl. At position 527, aspartic acid is changed to glutamic acid (527D>E), and at position 532, lysine is changed to glutamic acid (532K>E), preventing cleavage by trypsin. In our results, two peptides overlapped with this sequence, one featuring a missed cleavage, supported by 13 and 10 spectra, respectively. Figure 4D and 4E display two examples of highly confident matches, and Supplementary Table 1 lists PEP, q-value, and agreement with predictors for all PSMs. Altogether, the PEPs and agreement with predictors for these PSMs support the identification of this sequence, and thus the presence of these haplotypes in the data reported by Wang et al. (17), consistent with the frequencies of these haplotypes in the European population. The sequence encoded by these haplotypes cannot be detected using canonical databases.

While including the sequences from different haplotypes offers the ability to detect new protein haplotypes, it also increases the likelihood of similar peptides to map to different proteins. For example, the protein *POTE ankyrin domain family member I* (POTEI, ENSP00000392718) contains in its most frequent haplotype eight amino acid substitutions, two of which fall into the same tryptic peptide. In the actin-like domain of POTEI at position 918, tyrosine changes to phenylalanine (918Y>F), and at position 929 methionine changes into threonine (929M>T), thus encoding the peptide LC**F**VALDFEQEMA**T**AASSSSLEK (Table 1). The frequency of this haplotype among participants of the European population in the 1000 Genomes project is 46 %, while in this population the aggregated frequency of all haplotypes not containing any of these substitutions is 1.98 %. However, the sequence of the corresponding region of POTEI is highly similar to the sequence of actin beta (ACTB), actin gamma 1 (ACTG1), and actin alpha 1 (ACTA1), differing in 1, 1, and 2 residues, respectively. Such highly similar sequences represent peptides differing in their composition by only a few atoms, a mass difference that can be indistinguishable from a chemical or post-translational modification (e.g., a chemical modification of methionine can be mistaken for a substitution of methionine to threonine (29)). Therefore, telling these two proteins apart can be extremely challenging when accounting for variants. Conversely, if only canonical sequences are included in the database, the spectra obtained from POTEI will be arbitrarily assigned to actin. Numbers of spectra matching the corresponding regions of these proteins are listed in Table 1. Matching spectra to each of the peptide sequences in Table 1 have been identified in all three samples.

**Table 1:**
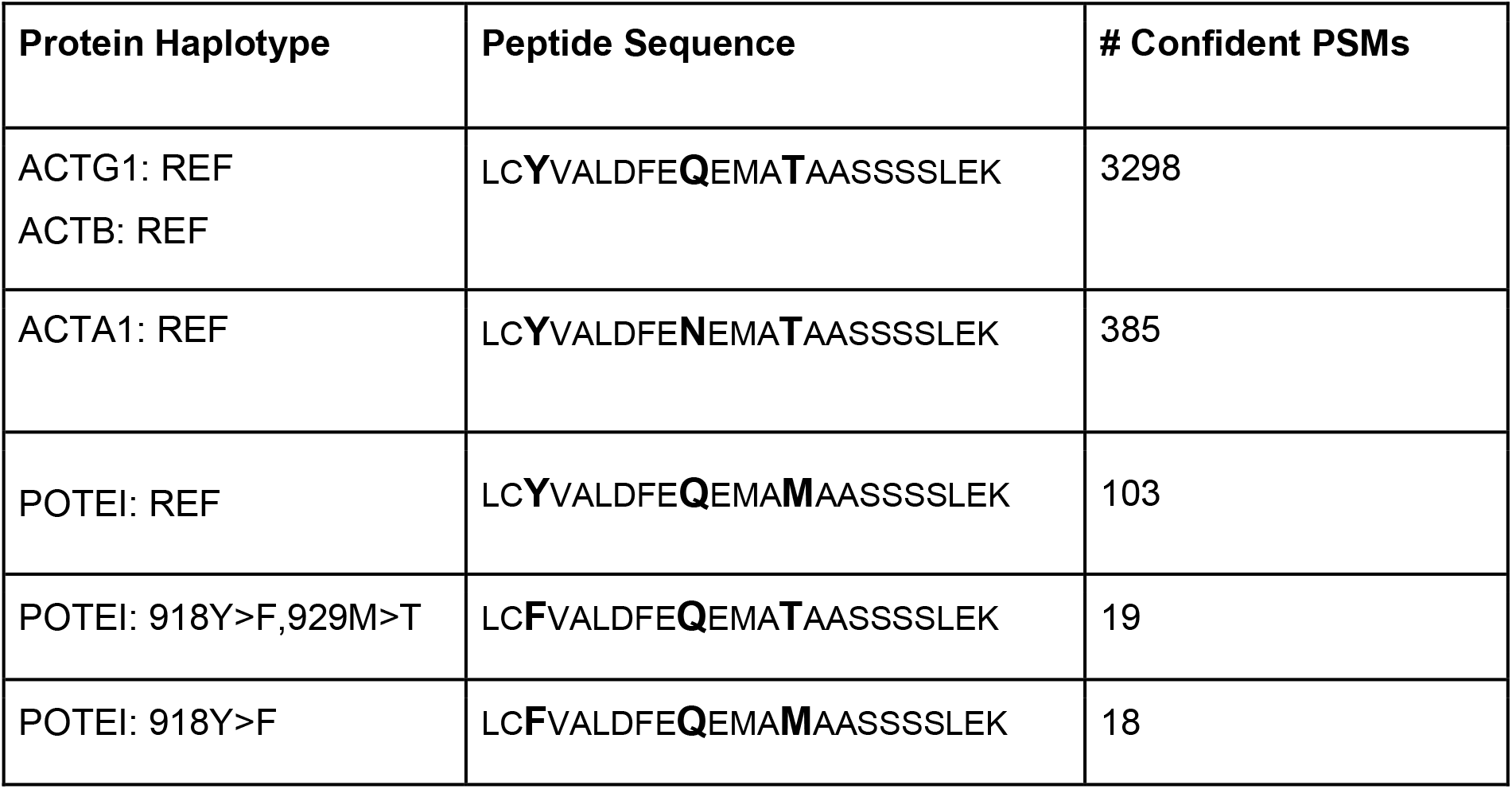
PSMs mapping to the five highly similar protein sequences: actin gamma 1 (ACTG1) / actin beta (ACTB), actin alpha 1 (ACTA1), and three haplotypes of POTEI. *REF* marks the canonical sequence. We specify the number of confident PSMs matching the sequence of interest, and number of samples with any spectra matching to these peptides.

| Protein Haplotype | Peptide Sequence | # Confident PSMs |
| --- | --- | --- |
| ACTG1: REF<br>ACTB: REF | LC**Y**VALDFE**Q**EMAT**A**ASSSSLEK | 3298 |
| ACTA1: REF | LC**Y**VALDFE**N**EMAT**A**ASSSSLEK | 385 |
| POTEI: REF | LC**Y**VALDFE**Q**EMAM**A**ASSSSLEK | 103 |
| POTEI: 918Y>F,929M>T | LC**F**VALDFE**Q**EMAT**A**ASSSSLEK | 19 |
| POTEI: 918Y>F | LC**F**VALDFE**Q**EMAM**A**ASSSSLEK | 18 |

The need to distinguish very similar sequences makes the use of haplotype-specific databases particularly sensitive to the spectrum identification strategy. As an example, we conducted the search again after enabling the refinement procedure of X!Tandem. This procedure is a multi-step approach that selects a limited set of proteins for a secondary search with different search parameters including more modifications and relaxing thresholds, e.g., in terms of missed cleavages. While this procedure presents the advantage to quickly scan for new peptides, it makes the evaluation of matches challenging (30), and increases the likelihood to encounter cases where a modification can be mistaken for an amino acid substitution, and vice versa. Figure 5 shows such an example of two matches to the same spectrum, obtained using the refinement procedure: one peptide contains the product of the alternative allele of two variants (Figure 5A) while the other has the product of the reference allele for one of the variants with a modification on the N-terminus compensating the mass difference (Figure 5B). Both matches show a good matching of the higher-mass peaks and good agreements with the predictors, but a high prevalence of unmatched peaks. Based on their scores, both matches would pass a 1 % FDR threshold, but the similarity between the sequences makes it challenging to assess whether one or the other haplotype is a better match. This example shows the difficulty to distinguish variant peptides when the amino acid substitution has a mass difference equal or very similar to a modification. Overall, we observed inflated identification rates for multi-variant peptides using the refinement procedure (1,060 PSMs with refinement vs. 342 PSMs without). For example, without the refinement procedure, 19 spectra matched the multi-variant sequence of POTEI among the PSMs passing a 1 % FDR threshold (Table 1); with the refinement procedure, the results contained 113 matching spectra. From the 94 additional matches, we suspect that many correspond to other sequences that were artifactually matched to this sequence, possibly through the addition of modifications.

**Figure 5:**
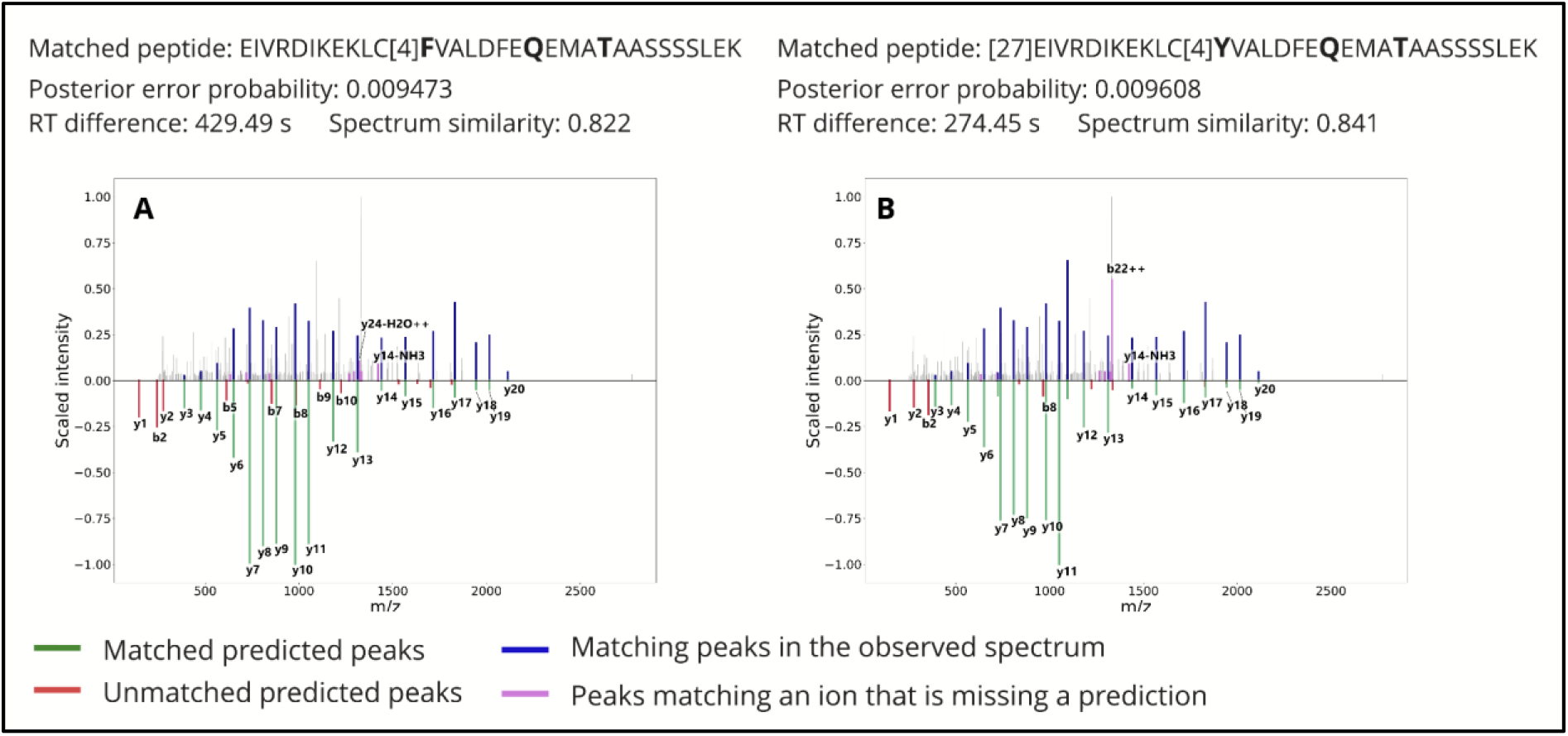
Comparison between predicted spectra for two different peptides matched to the same observed spectrum. The posterior error probability as obtained from Percolator is listed along with retention time difference to prediction as obtained from DeepLC, and spectrum similarity with prediction as obtained from MS2PIP. The intensity of the measured spectrum is plotted (top; blue, pink, and gray) with the scaled predicted intensity mirrored (bottom; green and red). Peaks in the measured spectrum matching predictions are highlighted in blue, measured peaks matching an ion with a missing intensity prediction are highlighted in pink, while other measured peaks are plotted in gray. Numbers in the peptide sequence are identifiers of post-translational modifications in UniMod (31).

Error rates derived from the target-decoy strategy rely on the modeling of the null distribution of scores using random matches. Distinguishing a variant peptide from a modified one however requires telling apart two matches that are very similar, and both better than random. In such cases, it is expected that modeling the null distribution using random matches provides underestimated error rates, and additional quality control measures can be applied to assess the quality of the matches (32). We submitted the variant matches passing the target-decoy 1 % FDR threshold in the X!Tandem search without the refinement procedure to PepQuery, a targeted peptide search engine providing additional validation for variant peptides identified using proteomics (33). PepQuery found that a substantial share of the matches were low scoring or could also match another peptide (10 % and 11 % of the matches, respectively), and the prevalence of these matches decreased with the PEP (Figure 6A). Conversely, 47 % of the matches were labeled as confident and the prevalence of confident matches increased with the PEP. The remaining matches were labeled as possibly matching a modification not considered in the original search - a rare post-translational modification or an artifact introduced during sample preparation. Interestingly, the prevalence of such ambiguous matches was stable around 30 % across PEP bins. These results highlight the difficulty posed by modifications in the confident identification of variant peptides. In the case of highly similar expected spectra between a variant and a modified peptide, analysts need to rely on prior knowledge on the likelihood of finding a given allele or modification in the sample studied, or on the presence of diagnostic ions (Figure 7). In the example of Figure 7B, the detection of y29++ would advocate in favor of the variant peptide rather than the modified peptide, but this peak is of low intensity and therefore represents only thin evidence.

**Figure 6:**
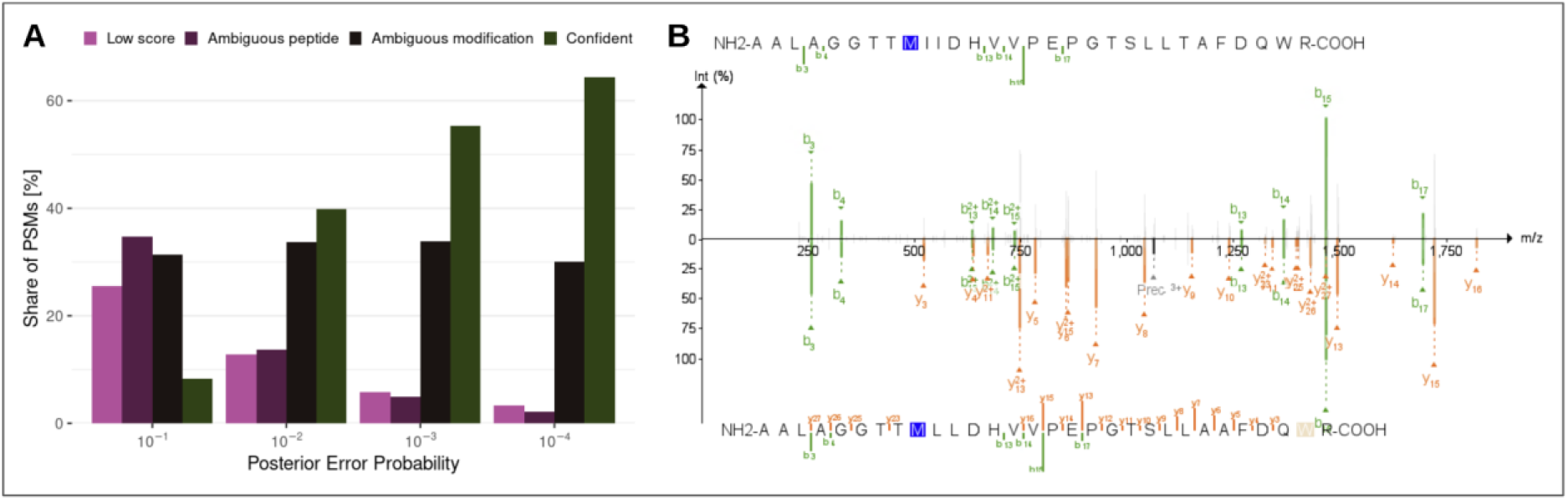
Analysis of variant peptides passing a target-decoy 1 % FDR threshold using PepQuery. A: histogram of the PSM type according to PepQuery. Low score: the match was not further investigated by PepQuery due to low score; Ambiguous peptide: the spectrum could be matched to a reference peptide at similar score; Ambiguous modification: the spectrum could be matched to a reference peptide at similar score when accounting for a modification that was not included in the original search; Confident: the match passed all PepQuery validation filters. B: Mirrored annotated spectra obtained using PDV (34) of a variant PSM with better match when accounting for a modification not included in the search, here a dioxydation of tryptophan.

**Figure 7:**
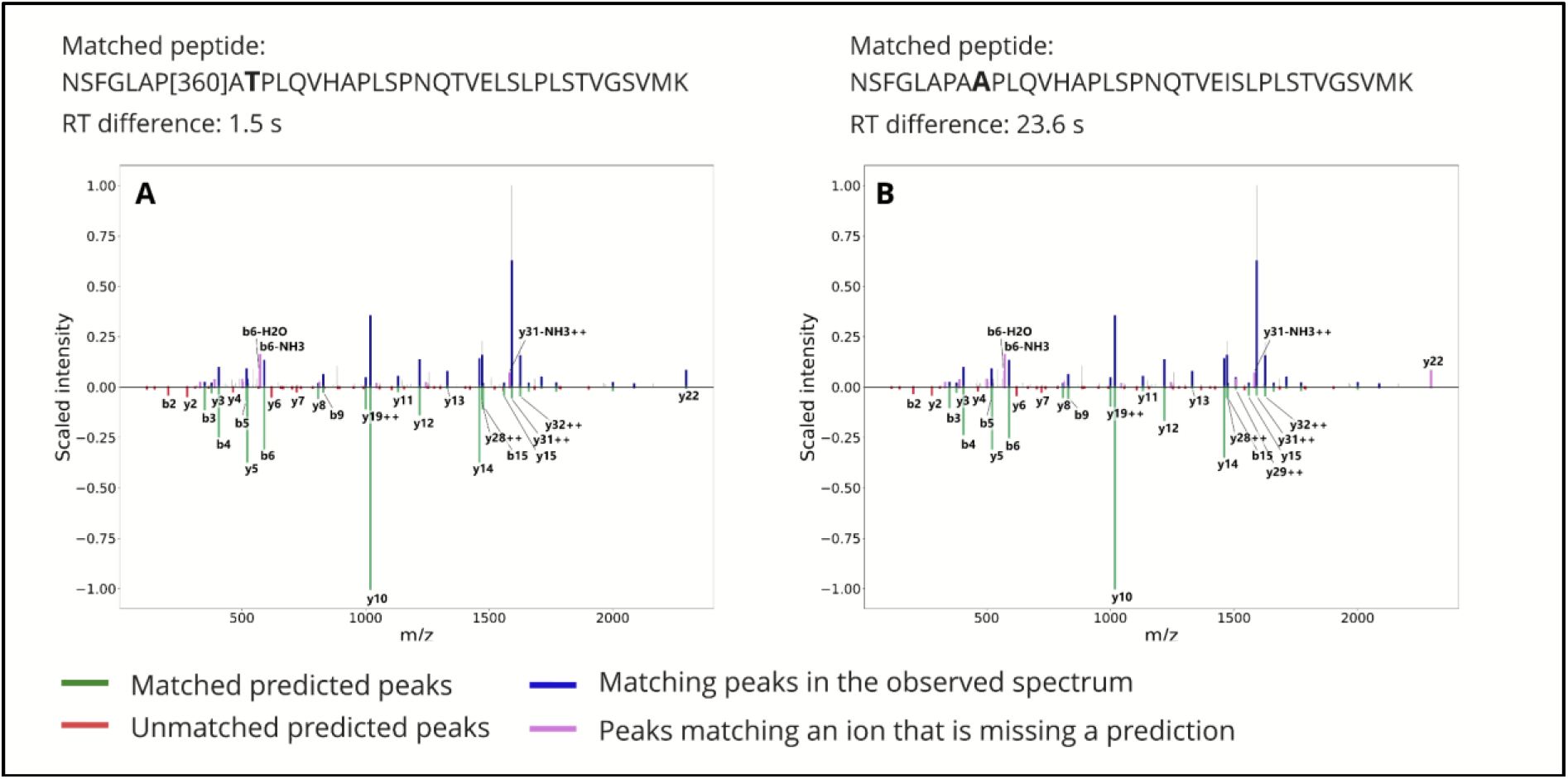
Comparison between predicted spectra as obtained from MS2PIP for two different peptides matched to the same observed spectrum. A: Peptide candidate suggested by PepQuery is a canonical sequence with a modification; B: Peptide candidate suggested in our search is a variant peptide. We list the retention time difference to prediction as obtained from DeepLC. The intensity of the measured spectrum is plotted (top; blue, pink, and gray) with the scaled predicted intensity mirrored (bottom; green and red). Peaks in the measured spectrum matching predictions are highlighted in blue, measured peaks matching an ion with a missing intensity prediction are highlighted in pink, while other measured peaks are plotted in gray. Numbers in the peptide sequence are identifiers of post-translational modifications in UniMod (31).

## Conclusion and Discussion

In this study, we propose to take advantage of the correlation between alleles through linkage disequilibrium to allow for the identification of peptides containing multiple linked amino acid substitutions, hence avoiding the computation of all possible combinations of alleles (14). Co-occurring alleles in the protein-coding regions of a gene yield specific protein sequences – protein haplotypes. Building upon previous work in proteogenomics, we created a search space of protein haplotypes. We observe that 7.69 % of the whole proteome maps to peptides that can contain an amino acid substitution, and up to 9.96 % of all discoverable substitutions are located in peptides where multiple substitutions co-occur (multi-variant peptides). These cases suggest that linkage disequilibrium between alleles resulting in amino acid substitutions should be included in a proteomics search space when identifying common variation. Subsequently, we performed a re-analysis of three samples of healthy tonsil tissue provided by Wang et al. (17). We identified peptides encoded by haplotypes containing 4,251 unique amino acid substitutions compared to the reference sequences of Ensembl, 6.63 % of which were found only in multi-variant peptides.

Although searching haplotype-specific sequences allows for the discovery of novel peptides that match to protein haplotypes, numerous challenges still remain. 57.9 % of the predicted haplotypes contain only substitutions, the remaining haplotypes contain other types of polymorphisms (insertions, deletions, or polymorphisms introducing or removing a stop codon). These cannot be detected using the sequences obtained from Haplosaurus. Moreover, with the introduction of haplotypes, the search space consists of a large number of proteoforms with a high degree of similarity, making it challenging to infer which proteoform has been identified. Amino acid substitutions of a mass difference equal to a post-translational or chemical modification are particularly challenging, as their distinction relies on the detection of few specific ions. This implies that searching without the correct haplotype or modification will generate incorrect sequences or modifications that are not caught by current error rate estimation strategies. Even worse, using the wrong haplotype on a protein sequence can result in a match in another protein. The prevalence of such errors in published proteomic datasets is currently unknown.

The dataset of protein haplotypes provided by Spooner et al. (1) was created using the genome assembly version GRCh37, which is now deprecated by Ensembl. During PepQuery analysis, we noted that a substantial share of variant peptides in GRCh37 would be canonical in GRCh38. For results that are fully up to date, a re-analysis of the data provided by the 1000 Genomes project on the current genome assembly is necessary. Limitations also come with the dataset of phased genotypes, as phasing may be inaccurate in regions with low linkage disequilibrium or in repetitive regions, resulting in an overestimation of haplotype frequencies (1). Finally, the methods for the scoring of confidence of peptide-spectrum matches are not well suited to distinguishing between multiple candidate sequences with a high degree of similarity. In the literature, the identification of variant peptides is validated by generating reference spectra using synthetic peptides (35,36), but such an approach presents a substantial cost and low throughput. In the present work, we used retention time and fragmentation predictors to generate the reference spectra *in silico*, and used these to evaluate the matches. Predictors can instead be directly coupled to Percolator, as implemented in MS2Rescore (37), and hence provide features that can improve the discrimination power between very similar peptides.

In conclusion, accounting for protein haplotypes in the search space for mass spectrometry-based proteomic identification improves the ability to cover relevant regions of the proteome, and holds the potential to be utilized in the medical context, given that the database of protein haplotypes is complete and up to date, and novel methods of quality control are developed.

## Acknowledgements

This work was supported by the Research Council of Norway (project #301178 to MV), the University of Bergen, and the Novo Nordisk Foundation (project NNF20OC0063872 to SJ).

The computations were performed on the Norwegian Research and Education Cloud (NREC), using resources provided by the University of Bergen and the University of Oslo. https://www.nrec.no

This research was funded, in whole or in part, by the Research Council of Norway 301178. A CC BY or equivalent license is applied to any Author Accepted Manuscript (AAM) version arising from this submission, in accordance with the grant’s open access conditions.

## Materials and Methods

### Database of protein sequences

The sequence database used for the search was built using data provided by Spooner et al. (1), who generated a database of protein haplotypes using their tool Haplosaurus. The haplotypes were generated using phased genotype data from the 1000 Genomes project Phase 3, obtained using methods described in (2). The haplotype analysis was performed using the transcript database Ensembl version 83 (38), human reference genome assembly version GRCh37 (1). The data provided by Spooner et al. (1) can be found at [https://doi.org/10.6084/m9.figshare.5545084]. For this work, we selected only protein haplotypes generated from alleles with FoO at least 1 % worldwide. This database was appended with the list of canonical protein sequences in the corresponding version of Ensembl, and a list of common sample contaminants, obtained from [https://www.thegpm.org/crap/]. The resulting search space contains 104,736 reference sequences, assembly version GRCh37, 290,080 protein haplotype sequences obtained as described above, and 116 sequences of sample contaminants. 394,959 decoy sequences were generated using the algorithm DecoyPyrat (39), provided by the tool *py-pgatk* (12). The final protein sequence database in the FASTA format is available as Supplementary material (SD2).

### Classification of peptides

We classified peptide sequences as canonical, single-variant, or multi-variant based on the number of amino acid substitutions they contain. If a peptide is canonical with respect to one protein sequence, and single-variant or multi-variant with respect to another protein sequence, it is classified as canonical. Similarly, if a peptide is a single-variant peptide with respect to one protein sequence, and multi-variant with respect to another protein sequence, it is classified as a single-variant peptide. Substitutions mapping to a peptide that has been “downgraded” in such manner are not considered as discovered, or discoverable.

### Public data reanalysis

We used this database to perform a reanalysis on a subset of data published and initially analyzed by Wang et al. (17) – 108 fractions from three samples of healthy tonsil tissue digested by trypsin, fragmented using higher-energy collisional dissociation (HCD) (MS experiment IDs: P013107, P010694, P010747).

The search was performed using the command-line interface of SearchGUI (40), employing the X!Tandem search algorithm (19), allowing for the oxidation of methionine as a variable modification, and carbamidomethylation of cysteine as a fixed modification. PeptideShaker (41) was used for post-processing of the search results and export of the PSMs to Percolator (20), which was used to evaluate the confidence of the matches and threshold using a false discovery rate (FDR) analysis (42). The list of PSMs was filtered to retain matches with a q-value below 0.01 (i.e., FDR is lower than 1 %). If a peptide matched to a contaminant sequence, it was removed from further analysis. As some of the canonical protein sequences in Ensembl contain multiple stop codons, the stop codon symbols were removed from their sequences for compatibility with X!Tandem. Peptides that would contain a stop codon were removed from further analysis.

### Quality control

To provide supporting evidence for the confidence of the peptide-spectrum matches (PSMs), chromatographic retention times were predicted by DeepLC (21), and expected peptide fragment ion intensities were predicted using MS2PIP (22). Peptides passing the 1 % FDR threshold were used for calibration of the DeepLC predictions. The absolute distance between the centered and scaled predicted and observed retention times was computed. The MS2PIP predictions were used to measure the distance between the predicted and observed spectrum. The peaks are scaled so that the median intensity in the observed spectrum corresponds to the median intensity in the prediction. A peak in the observed spectrum is considered matching to a peak in the prediction if it differs in m/z by no more than 10 ppm.

The distance between the matched predicted peaks and the observed ones is expressed as their angular similarity, calculated by the formula in equations 1 and 2:

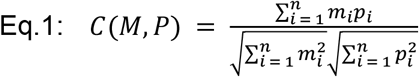

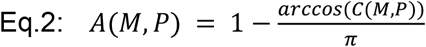

Where *M*=*(m_1_,…,m_n_)* is the set of intensities for the matched measured peaks, and *P*=*(p_1_,…,p_n_)* is the set of intensities for the matched predicted peaks, *n* is the number of matched peaks in the spectrum. *C(M,P)* denotes the cosine similarity between *M* and *P*, and *A(M,P)* denotes the angular similarity between *M* and *P*.

Predicted and observed spectra were also displayed as mirror plots for visual comparison in selected PSMs. The peaks in the observed spectrum matching to a predicted peak are highlighted in blue. As the intensity prediction for certain ion fragments by MS2PIP is missing, peaks matching those ions are highlighted in pink. The remaining measured peaks are displayed in gray. Peaks of the predicted spectrum are shown as negative values and labeled by the corresponding fragment ion. The predicted peaks that match a measured peak are displayed in green, unmatched predicted peaks are displayed in red.

### PepQuery Analysis

The variant PSMs passing 1 % FDR at PSM level using X!Tandem were further validated using PepQuery (v2.0.3) (33). The following parameters were used: Fixed modifications, Carbamidomethylation of C; variable modifications, Oxidation of M, Ammonia-loss of C, Glu->pyro-Glu of E, Gln->pyro-Glu of Q, Acetylation of peptide N-term; Precursor ion mass tolerance, 20 ppm; MS/MS mass tolerance, 0.05 Da; Enzyme specificity, trypsin; maximum missed cleavages, 2; allowed isotope range: −1,0,1,2. The parameter “-hc” was also set in the analysis. The human protein database from GENCODE Release 43 (GRCh37) was used as the reference protein database in the validation. The PSMs that passed all the filtering steps in PepQuery were considered confident. A complete list of variant PSMs with possible alternative peptide candidates suggested by PepQuery is available as Supplementary material (SD5).

## Data Availability

Supplementary data can be downloaded from [https://doi.org/10.6084/m9.figshare.21408117]. We provide the following files:

SD1: FASTA file including all target protein sequences (Ensembl reference proteome, protein haplotype sequences, contaminant sequences), excluding decoys

SD2: FASTA file including all target and decoy sequences

SD3: List of all peptide-spectrum matches (PSMs), resulting from the first run of X!Tandem without the refinement procedure, with all related metadata and quality control measures

SD4: List of substitutions identified, along with IDs of corresponding PSMs

SD5: List of variant PSMs and peptide candidates suggested by PepQuery, along with confidence scores for each peptide candidate

The pipeline to reproduce the post-processing steps, as well as a further description of the resulting files, are provided in https://github.com/ProGenNo/IdentifyingHaplotypesByMS.

## Abbreviations

FDR: false discovery rate
HCD: higher-energy collisional dissociation
LC: liquid chromatography
LD: linkage disequilibrium
MS: mass spectrometry
MS/MS: tandem mass spectrometry
PEP: posterior error probability
PSM: peptide-spectrum match
PTM: post-translational modification

---

**Supplementary Table 1:** PSMs matching to the multi-variant peptide covering a region of the most common haplotype of the IGQAP2 protein, and their respective confidence measures. The posterior error probability and q-value as obtained from Percolator are listed along with retention time difference to prediction as obtained from DeepLC, and spectrum similarity with prediction as obtained from MS2PIP.

| Sequence | RT error | Spectra angular similarity | q-value | Posterior error prob. |
| --- | --- | --- | --- | --- |
| VLWLDEIQQAVDEANVDEDR | 320.75 | 0.786 | 0.000113 | 0.000088 |
| VLWLDEIQQAVDEANVDEDR | 417.39 | 0.800 | 0.000113 | 0.000109 |
| VLWLDEIQQAVDEANVDEDR | 320.75 | 0.795 | 0.000113 | 0.000126 |
| VLWLDEIQQAVDEANVDEDR | 394.89 | 0.793 | 0.000129 | 0.000161 |
| VLWLDEIQQAVDEANVDEDRAK | 425.21 | 0.843 | 0.000150 | 0.000173 |
| VLWLDEIQQAVDEANVDEDR | 320.75 | 0.741 | 0.000150 | 0.000174 |
| VLWLDEIQQAVDEANVDEDRAK | 620.04 | 0.900 | 0.000158 | 0.000179 |
| VLWLDEIQQAVDEANVDEDR | 394.89 | 0.794 | 0.000168 | 0.000196 |
| VLWLDEIQQAVDEANVDEDRAK | 425.21 | 0.829 | 0.000170 | 0.000198 |
| VLWLDEIQQAVDEANVDEDRAK | 620.04 | 0.876 | 0.000171 | 0.000213 |
| VLWLDEIQQAVDEANVDEDRAK | 407.30 | 0.835 | 0.000176 | 0.000232 |
| VLWLDEIQQAVDEANVDEDR | 417.39 | 0.778 | 0.000179 | 0.000266 |
| VLWLDEIQQAVDEANVDEDRAK | 428.68 | 0.688 | 0.000179 | 0.000268 |
| VLWLDEIQQAVDEANVDEDR | 430.84 | 0.752 | 0.000179 | 0.000268 |
| VLWLDEIQQAVDEANVDEDRAK | 598.31 | 0.856 | 0.000179 | 0.000270 |
| VLWLDEIQQAVDEANVDEDR | 417.39 | 0.802 | 0.000179 | 0.000289 |
| VLWLDEIQQAVDEANVDEDR | 320.75 | 0.732 | 0.000179 | 0.000297 |
| VLWLDEIQQAVDEANVDEDRAK | 425.21 | 0.669 | 0.000215 | 0.000398 |
| VLWLDEIQQAVDEANVDEDR | 394.89 | 0.788 | 0.000223 | 0.000411 |
| VLWLDEIQQAVDEANVDEDRAK | 620.04 | 0.739 | 0.000252 | 0.000519 |
| VLWLDEIQQAVDEANVDEDRAK | 620.04 | 0.713 | 0.000335 | 0.000758 |
| VLWLDEIQQAVDEANVDEDR | 430.84 | 0.807 | 0.000345 | 0.000812 |
| VLWLDEIQQAVDEANVDEDR | 320.75 | 0.772 | 0.000704 | 0.002311 |

